# Surveillance based dynamic empirical therapy in a health care facility: an artificial intelligence approach

**DOI:** 10.1101/611210

**Authors:** Nicolas Houy, Julien Flaig

## Abstract

We present a solution method to the problem of choosing empirical treatments that minimize the cumulative infected patient-days in the long run in a health care facility. We rely on the stochastic version of a compartmental model to describe the spread of an infecting organism in the health care facility, and the emergence and spread of resistance to two drugs. We assume that the parameters of the model are known. Empirical treatments are chosen at the beginning of each period based on the count of patients with each health status. The same treatment is then administered to all patients, including uninfected patients, during the period and cannot be adjusted until the next period. Our solution method is a variant of the Monte-Carlo tree search algorithm. In our simulations, it allows to reduce the average cumulative infected patient-days over two years by 47.0% compared to the best standard therapy. We explain how our algorithm can be used either to perform online optimization, or to produce data for quantitative analysis.

## 1 Introduction

Health care facilities are particularly affected by infections with drug resistant pathogens [5]. There are two main reasons for this. First, the spread of infections (whether drug resistant or not) can be easier in hospital environments [9] and hospital patients are more at risk of serious complications than the general population. Second, antimicrobials are heavily used in health care facilities, both for preventive reasons and empirically. Empirical therapies are antimicrobial treatments administered at the onset of symptoms, when a therapy needs to be started immediately, before test results are available. But under antimicrobial treatment, drug resistant mutant strains are favored as compared to drug susceptible strains. Antimicrobials select for resistance both at the within-host and between-host levels [12].

Empirical prescription policies have been proposed to slow and even revert the evolution of resistance [14]. Indeed, at the molecular level, resistance is acquired through the modification of essential pathways of the organism. Therefore, it is often^1^ associated with fitness costs: in the absence of treatment, resistant organisms are less able to survive and reproduce than wild-type susceptible organisms [1, 13]. It was then hoped that empirical therapy policies such as drug mixing – assigning drugs randomly to patients – or drug cycling – switching from one drug to another following a rotation schedule – would limit the emergence and spread of antimicrobial resistance. Yet the respective merits of drug mixing and drug cycling have been debated, and empirical studies remain inconclusive (see [2–4, 11, 15, 16, 18] and the references therein).

The theoretical investigations of empirical therapy policies typically rely on aggregate (between-host) models of the emergence and spread of antimicrobial resistance; see [17] for a review. In this line, it was shown in [10] that policies based on the results of microbiological tests could limit the spread of resistance. In this scenario, all symptomatic patients are tested at a given rate in order to choose an adequate individual treatment. Asymptomatic patients and not yet tested symptomatic patients receive an empirical treatment based on available test results. A significant advantage of this strategy is that it makes use of information available at no additional cost since tests are performed by default for all symptomatic patients.

In this article, we approach the problem of devising empirical therapies based on microbiological test results from another angle. First, we strive for *optimal* policies. By contrast, the policies presented in [10] are *adaptive* in that they specify which empirical treatment is to be administered given test results that depend only on the spread of the disease in the past and on past treatments. Of course, the merit of an empirical therapy at a given time depends not only on past treatments, but also on the subsequent course of the epidemic that is on subsequent empirical treatments. We solve this dynamic optimization problem with a variant of the Monte-Carlo tree search algorithm, a solution method borrowed from the field of artificial intelligence. See [6] for an introductory application of this method to empirical therapy optimization, and [7, 8] for an application to chemotherapy regimen optimization. As for the underlying epidemiological model, our optimization method relies on the stochastic version of a compartmental model derived from those presented in [17].

Second, we assume that we periodically have access to the health status of all patients in the hospital. That is, we are able to test all patients with a fixed period, and test results are available immediately. In [10], information becomes available continuously, but it is only partial – test results are used as a proxy for actual prevalence. Also, we assume that all patients, including uninfected patients,^2^ receive the same treatment; and that treatments are not continuously adjusted dynamically during periods between observations. Notice finally that since our empirical therapies that are contingent on the exact number of patients with each health status, relying on stochastic models is necessary.

We present the materials and methods used in the study in Section 2: the compartmental model describing the emergence and spread of a microbial disease with resistant strains in a health care facility in Section 2.1, and the optimization algorithm in Section 2.2. We show and discus our results in Section 3. Section 4 concludes.

## 2 Materials and methods

### 2.1 Model

A sketch of our epidemiological model is shown in Figure 1. We summarize the considered events in Table 1 and the parameters in Table 2. A detailed description and discussion of the model can be found in [6].

**Table 1:** Events considered in the model.

| Population changes | Rate |
| --- | --- |
| *Admissions and discharges* |  |
| $S \leftarrow S + 1$ | $N\mu m_S$ |
| $R_i \leftarrow R_i + 1, i \in \{1, 2, 12\}$ | $N\mu m_i$ |
| $X \leftarrow X + 1$ | $N\mu \left(1 - \sum_{i \in \{S, 1, 2, 12\}} m_i\right)$ |
| $S \leftarrow S - 1$ | $\mu S$ |
| $R_i \leftarrow R_i - 1, i \in \{1, 2, 12\}$ | $\mu R_i$ |
| $X \leftarrow X - 1$ | $\mu X$ |
| *De novo mutations* |  |
| $S \leftarrow S - 1, R_i \leftarrow R_i + 1, i \in \{1, 2\}$ | $\nu_i [f_0 \chi_\epsilon(s_i, s_S) + f_i \chi_\epsilon(s_i, 1)] S$ |
| $S \leftarrow S - 1, R_{12} \leftarrow R_{12} + 1$ | $\nu_{12} [f_0 \chi_\epsilon(s_{12}, s_S) + (f_1 + f_2 + f_{12}) \chi_\epsilon(s_{12}, 1)] S$ |
| $R_i \leftarrow R_i - 1, R_{12} \leftarrow R_{12} + 1, i \in \{1, 2\}$ | $\nu_{(3-i)} [(f_0 + f_i) \chi_\epsilon(s_{12}, s_i) + (f_{(3-i)} + f_{12}) \chi_\epsilon(s_{12}, 1)] R_i$ |
| *Infections and recoveries* |  |
| $X \leftarrow X - 1, S \leftarrow S + 1$ | $f_0 \beta (1 - c_S) \chi_\epsilon(s_S, 1) X S / N$ |
| $X \leftarrow X - 1, R_i \leftarrow R_i + 1, i \in \{1, 2\}$ | $(f_0 + f_{(3-i)}) \beta (1 - c_i) \chi_\epsilon(s_i, 1) X R_i / N$ |
| $X \leftarrow X - 1, R_{12} \leftarrow R_{12} + 1$ | $\beta (1 - c_{12}) \chi_\epsilon(s_{12}, 1) X R_{12} / N$ |
| $S \leftarrow S - 1, X \leftarrow X + 1$ | $[\tau(f_1 + f_2 + f_{12}) + \gamma] S$ |
| $R_i \leftarrow R_i - 1, X \leftarrow X + 1, i \in \{1, 2\}$ | $[\tau(f_{(3-i)} + f_{12}) + \gamma] R_i$ |
| $R_{12} \leftarrow R_{12} - 1, X \leftarrow X + 1$ | $\gamma R_{12}$ |
| *Superinfections* |  |
| $S \leftarrow S - 1, R_i \leftarrow R_i + 1, i \in \{1, 2\}$ | $\sigma \beta (1 - c_i) [f_0 \chi_\epsilon(s_i, s_S) + f_i \chi_\epsilon(s_i, 1)] S R_i / N$ |
| $R_i \leftarrow R_i - 1, S \leftarrow S + 1, i \in \{1, 2\}$ | $\sigma \beta (1 - c_S) f_0 \chi_\epsilon(s_S, s_i) S R_i / N$ |
| $S \leftarrow S - 1, R_{12} \leftarrow R_{12} + 1$ | $\sigma \beta (1 - c_{12}) [f_0 \chi_\epsilon(s_{12}, s_S) + (f_1 + f_2 + f_{12}) \chi_\epsilon(s_{12}, 1)] S R_{12} / N$ |
| $R_{12} \leftarrow R_{12} - 1, S \leftarrow S + 1$ | $\sigma \beta (1 - c_S) f_0 \chi_\epsilon(s_S, s_{12}) S R_{12} / N$ |
| $R_i \leftarrow R_i - 1, R_{12} \leftarrow R_{12} + 1, i \in \{1, 2\}$ | $\sigma \beta (1 - c_{12}) [(f_0 + f_i) \chi_\epsilon(s_{12}, s_i) + (f_{(3-i)} + f_{12}) \chi_\epsilon(s_{12}, 1)] R_i R_{12} / N$ |
| $R_{12} \leftarrow R_{12} - 1, R_i \leftarrow R_i + 1, i \in \{1, 2\}$ | $\sigma \beta (1 - c_i) (f_0 + f_i) \chi_\epsilon(s_i, s_{12}) R_i R_{12} / N$ |
| $R_i \leftarrow R_i - 1,$<br>$R_{(3-i)} \leftarrow R_{(3-i)} + 1, i \in \{1, 2\}$ | $\sigma \beta (1 - c_{(3-i)}) [f_0 \chi_\epsilon(s_{(3-i)}, s_i) + f_{(3-i)} \chi_\epsilon(s_{(3-i)}, 1)] R_i R_{(3-i)} / N$ |

**Table 2:** Model parameters.

| Parameter | Unit | Description | Value |
| --- | --- | --- | --- |
| *Admissions and discharges* |  |  |  |
| $N$ | – | Average number of patients. | 40 |
| $\mu$ | day<sup>-1</sup> | Turnover rate. | 0.1 |
| $m_S$ | – | Proportion of incoming patients infected with the susceptible strain. | 0.1 |
| $m_1$ | – | Proportion of incoming patients infected with the strain resistant to drug 1. | 0.05 |
| $m_2$ | – | Proportion of incoming patients infected with the strain resistant to drug 2. | 0.01 |
| $m_{12}$ | – | Proportion of incoming patients infected with the strain resistant to drugs 1 and 2. | 0.005 |
| *De novo mutations and fitness costs* |  |  |  |
| $\nu_i$ | day<sup>-1</sup> | Rate of acquisition of resistance to treatment $i$ through *de novo* mutation. $i \in \{1, 2, 12\}$ | 0.005 |
| $c_i$ | – | Cost on infectivity of resistance to treatment $i$, $i \in \{1, 2, 12\}$. | 0.2 |
| $c_S$ | – | Cost on infectivity of susceptible organisms. | 0 |
| $s_i$ | – | Cost on competitive performance of resistance to treatment $i$, $i \in \{1, 2, 12\}$. | 0.2 |
| $s_S$ | – | Cost on competitive performance of susceptible organisms. | 0 |
| $s_X$ | – | Cost on competitive performance of commensal organisms. | 1 |
| $\epsilon$ | – | Slope parameter in function $\chi_\epsilon$. | 0.5 |
| *Infections, superinfections, and recoveries* |  |  |  |
| $\beta$ | day<sup>-1</sup> | Transmission rate. | 0.5 |
| $\sigma$ | – | Relative rate of superinfection. | 1 |
| $\gamma$ | day<sup>-1</sup> | Rate of spontaneous recovery. | 0.1 |
| $\tau$ | day<sup>-1</sup> | Rate of recovery under appropriate treatment. | 0.3 |

**Figure 1:**
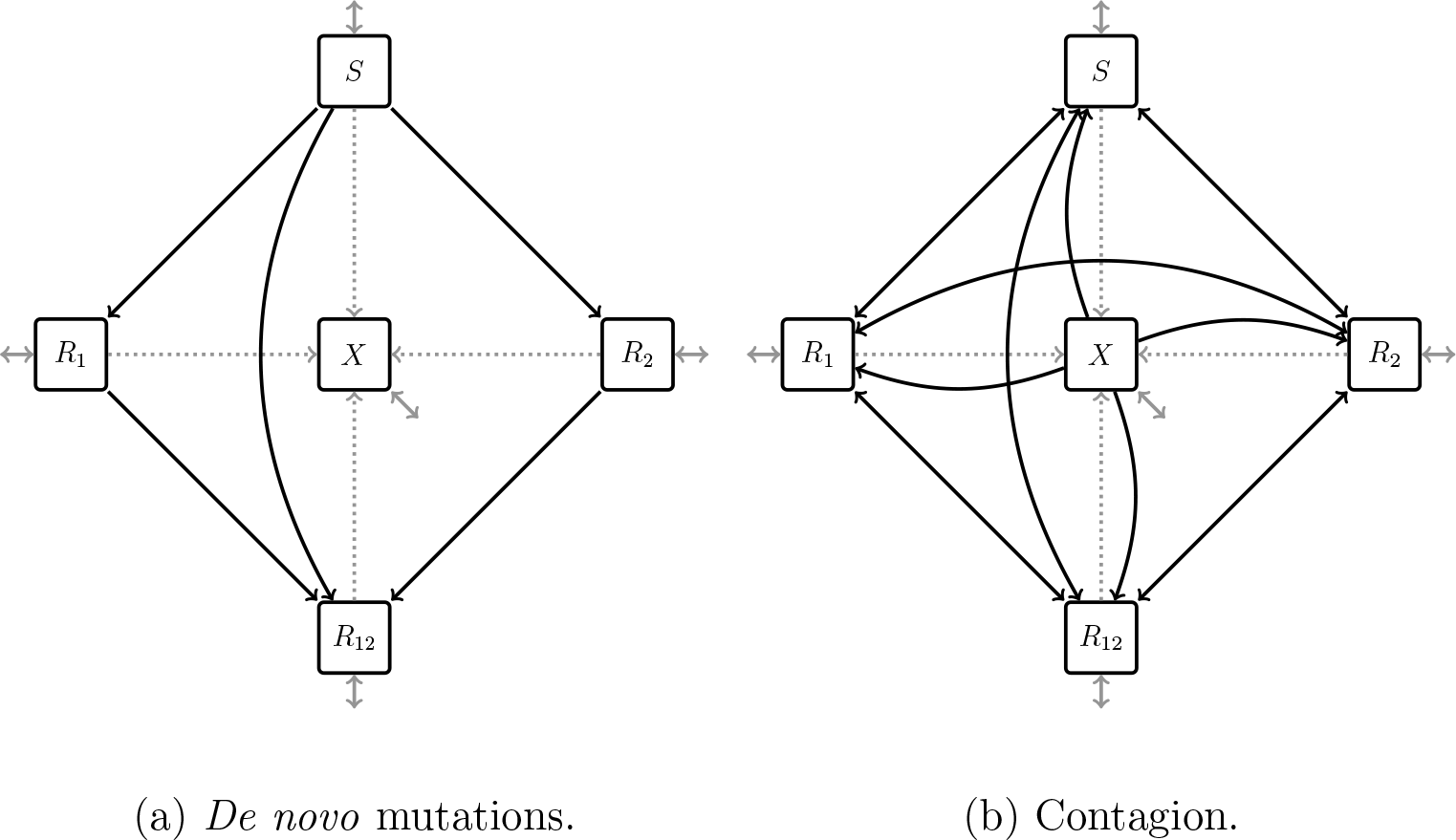
For clarity we display *de novo* mutations *(black, panel (a))* and contagion *(black, panel (b))* on two separate graphs. *Gray:* admissions and discharges. *Dotted gray:* recovery.

- *Pathogenic strains.* We model resistance to two drugs, drug 1 and drug 2. At each time, a patient is either infected with a wild-type strain susceptible to both drugs, infected with a strain resistant to drug 1 but not to drug 2 (1-resistant), infected with a strain resistant to drug 2 but not to drug 1 (2-resistant), infected with a strain resistant to both drugs (12-resistant), or uninfected with any of the above. *S*, *R*_1_, *R*_2_, *R*_12_, and *X* denote the number of patients with each health status respectively. We assume that infected patients are only infected with one pathogenic strain at any given time.
- *Admissions and discharges.* We consider a small health care facility or a hospital ward with an average population of *N* = 40 patients. We assume that admission and discharge do not depend on health status. However, we also make the simplifying assumption that transmission of strains and emergence of resistance in the community is exogenous to the problem in hand.
- *Treatments.* Three treatments are available: drug 1 in monotherapy (treatment 1), drug 2 in monotherapy (treatment 2), and a combination of drugs 1 and 2 (treatment 12). Patients can also be left without a treatment (treatment 0). Decision variable *f*_*i*_, *i* ∈ {0, 1, 2, 12}, is equal to 1 when patients receive treatment *i* and to 0 otherwise. At all time, *f*_0_ + *f*_1_ + *f*_2_ + *f*_12_ = 1.
- *Recovery.* Infected patients recover spontaneously at rate γ regardless of the infecting strain. Patients receiving adequate treatment recover at rate *τ*.
- De novo *mutation.* Infecting organisms acquire *i*-resistance through *de novo* mutation at rate *ν*_*i*_, *i* ∈ {1, 2, 12}, of two different genes as illustrated in Figure A.1 in Appendix.
- *Infection and superinfection.* We assume homogeneous mixing. Pathogenic strains are transmitted to uninfected patients with transmission rate *β*, and to already infected patients with transmission rate *σβ*. We assume *σ* = 1 in this instance of the model, but the general case is *σ* ∈ [0, 1]. *i*-resistant strains incur a fitness cost on transmission *c*_*i*_ ∈ [0, 1], *i* ∈ {1, 2, 12}, and their transmission rate is weighed by 1 − *c*_*i*_.
- *Strain replacement.* Within-host, a new pathogenic strain acquired through *de novo* mutation or contagion must compete with the resident pathogenic strain or with the commensal microflora in the case of previously uninfected patients. *i*-resistant strains incur a fitness cost *s*_*i*_ ∈ [0, 1], *i* ∈ {1, 2, 12}, on competitive performance. We normalize the fitness cost *s*_*S*_ of the wild-type susceptible strain to 0, and the fitness cost *s*_*X*_ of the commensal microflora to 1. A strain receiving adequate treatment has a fitness cost of 1. The rate of replacement of a resident strain with fitness cost *s*_*j*_ by a new strain with fitness cost *s*_*i*_ is given by function *χ*_*ϵ*_ as

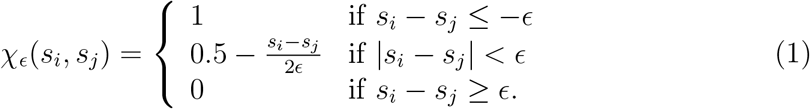

We further assume that a new strain cannot colonize a treated patient unless it is resistant to the treatment.

### 2.2 Optimization algorithm

Our objective is to minimize the cumulative infected patient-days in the long run (two years). We assume that we are able to know the exact number of patients with each health status every Δ*h* days. In the following, unless otherwise specified, Δ*h* = 15 days. Upon observation of patients’ health status, we must decide on a treatment to be administered empirically to all patients over the next Δ*h* days. This treatment cannot be changed over Δ*h* days. This precludes adjustments of the treatment administered to all patients, but also adjustments of individual treatments based for instance on the evolution of the symptoms.

Function STATETOTREATMENT (Algorithm 1) returns the treatment to be used over the next Δ*h* days given the observed state of the epidemic. Notice that our dynamic model is Markovian so that all the information regarding the past dynamics is indeed summarized with no loss in the current state of the epidemic. Θ denotes the set of available treatments among which we are to choose. It includes treatments 0, 1, 2, and 12. The treatment administered during the next Δ*h* days will of course have an influence on the subsequent spread of the disease, on the emergence of drug resistance, and therefore on the cumulative infected patient-days. However the long term effects of this treatment will also depend on subsequent empirical treatment decisions, which are yet unknown. Therefore, we decide on the treatment to be administered during the next Δ*h* days by assuming that a default treatment *θ*_*default*_ will be administered afterwards over an horizon *H* − Δ*h*, where *H* is equal to two years (720 days). Since we know the population dynamics parameter of the health care facility and of the disease, we are able to compare treatments by running stochastic simulations of the model presented in Section 2.1, starting from the observed state of the epidemic. For each available treatment *θ* ∈ Θ, we run *n*_*S*_ = 800 stochastic simulations assuming that *θ* is administered over Δ*h* days, and that default treatment *θ*_*default*_ is administered during the following *H* − Δ*h* days. We choose the treatment *θ*\* for which the average over *n*_*S*_ simulations of the cumulative infected patient-days over *H* is the smallest.

The results presented in this article were obtained using treatment 2 as default treatment. Notice that different default treatments will lead to different decisions. A default treatment must be chosen beforehand by simulating the use of the algorithm with different default treatments. Which default treatment performs better than the others will depend on the parameters of the model.

#### Algorithm 1 Choice of a policy given the observation of the number of patients in each health status

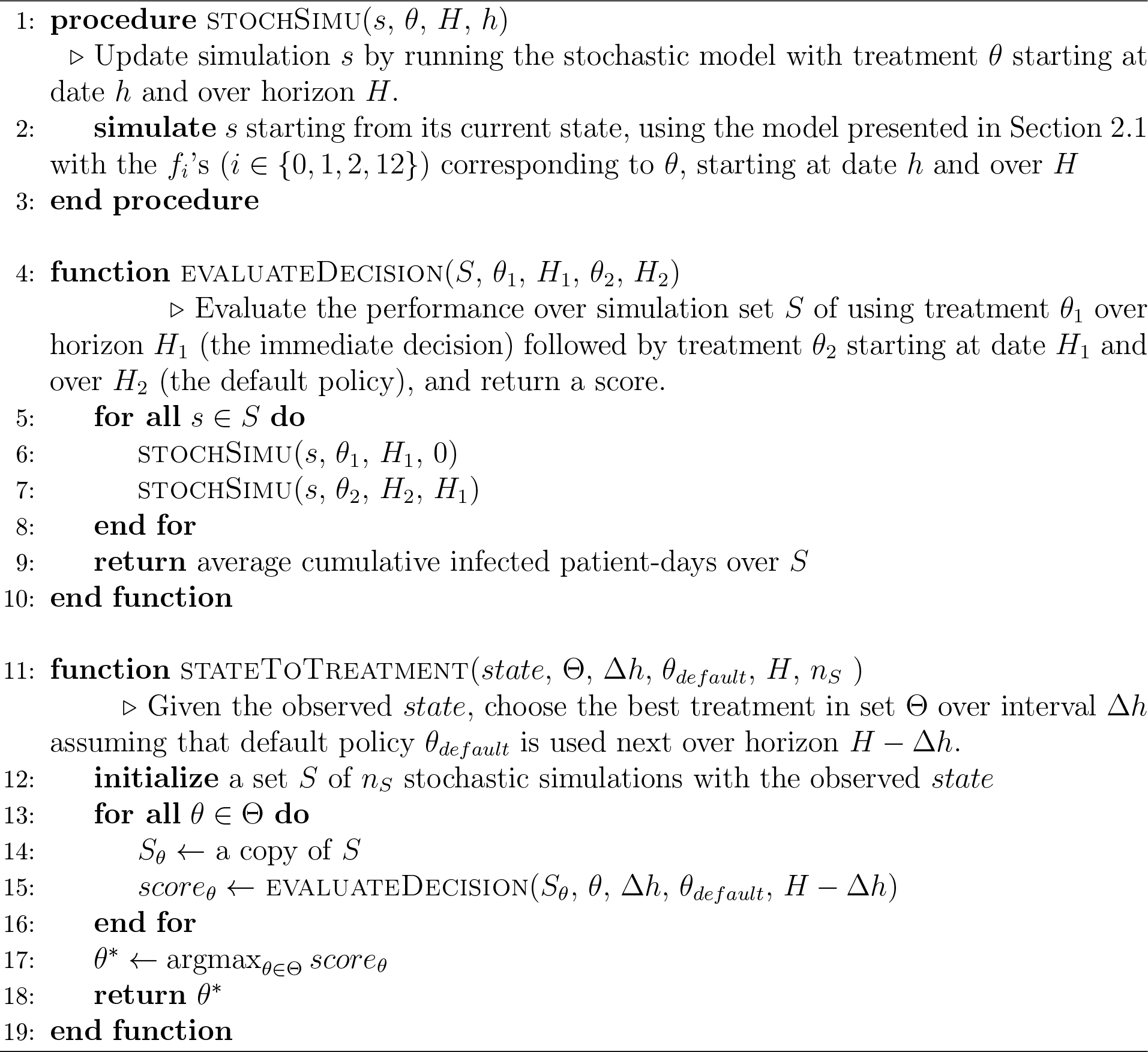

## 3 Results

We ran 400 simulations using the optimization algorithm presented in Section 2.2. Each simulation was initialized with a period of 30 years without treatment. A treatment was then chosen for each period of Δ*h* days over two years (720 days) by observing the number of patients at the beginning of the period and calling function stateToTreatment. Thus, we obtained 400 different empirical treatment regimens (shown in Figure A.2 in Appendix).

We call OPTIDYN the empirical therapy policy that consists in using our optimization algorithm to choose empirical treatments. In Figure 2, we compare the performance of OPTIDYN over two years with that of the following policies: NONE, that consists in administering no treatment; COMBO, that consists in administering treatment 12; CYC-30, that consists in alternating 30 days of treatment 1 with 30 days of treatment 2; MONO-1 that consists in treatment 1 in monotherapy; and MONO-2, that consists in treatment 2 in monotherapy. Finally, we compare OPTIDYN with OPTI, a policy that consists in choosing a treatment regimen for two years with drug switches allowed every 30 days, based on no other information than the population dynamics parameters, as explained in [6]. ^3^

**Figure 2:**
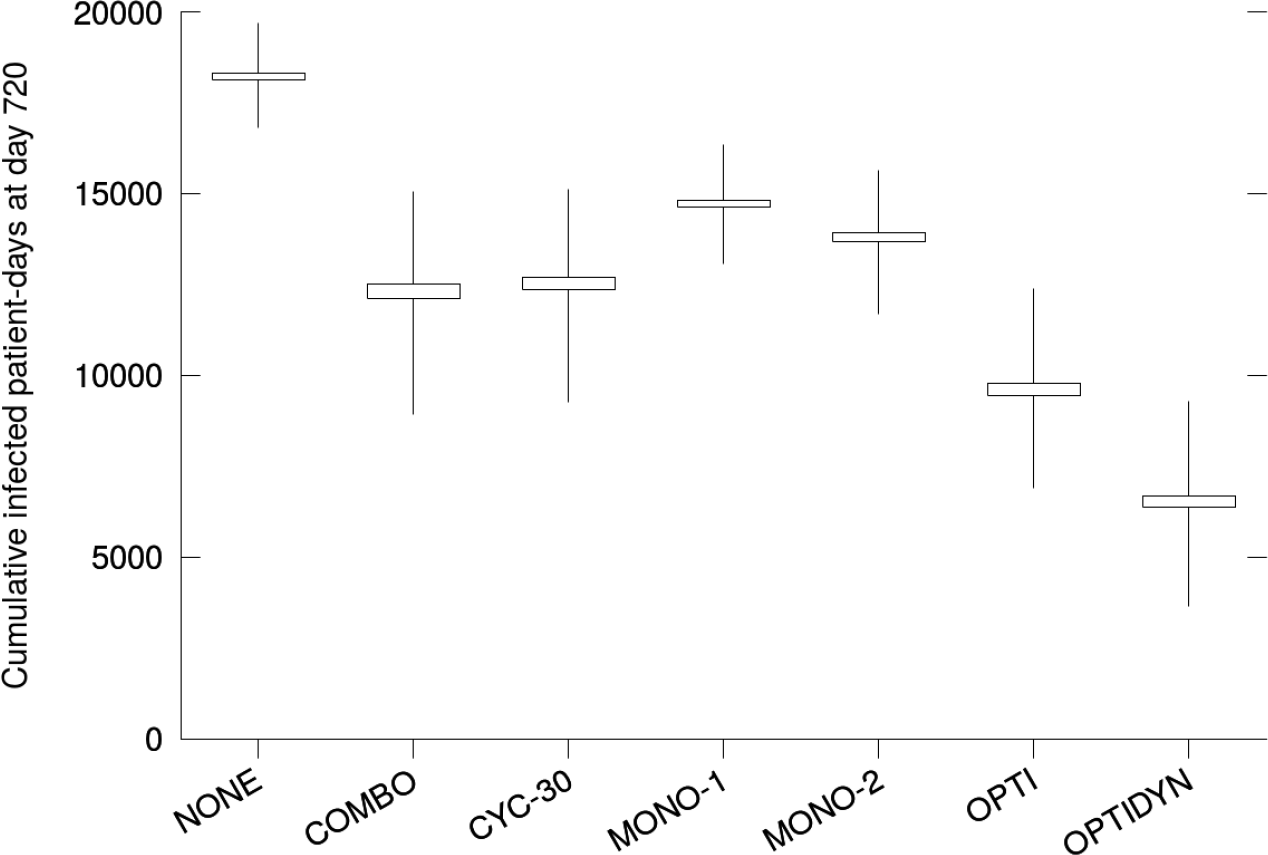
Box plot of the cumulative infected patient-days (400 simulations). *Boxes:* 95% confidence intervals. *Whiskers:* 5^th^ and 95^th^ percentiles of the simulations.

Without treatment (NONE), we obtain an average of 18,215.4 (95% CI: 18,126.2 – 18,304.6) cumulative infected patient-days over two years. The combination therapy COMBO and the cycling therapy CYC-30 show relatively close performance with an average of 12,314.5 (95% CI: 12,122.3 – 12,506.8) and 12,525.3 (95% CI: 12,355.1 – 12,695.6) cumulative infected patient-days respectively. The former allows a 32.4% and the latter a 31.2% reduction in average cumulative infected patient-days over two years compared to the NONE policy. COMBO and CYC-30 perform better than MONO-1 and MONO-2, which yield on average 14,716.7 (95% CI: 14,618.9 – 14,814.6) and 13,813.8 (95% CI: 13,689.7 – 13,937.9) cumulative infected patient-days. OPTI allows to decrease the average cumulative infected patient-days by 47.1% compared to NONE and by 21.8% compared to COMBO, to 9,629.61 (95% CI: 9,462.96 – 9,796.25) infected patient-days. With an average of 6,531.13 (95% CI: 6,365.9 – 6,696.36) cumulative infected patient-days over two years, OPTIDYN yields a reduction of 64.1% compared to NONE, of 47.0% compared to COMBO, and of 32.2% compared to OPTI (the three reductions are significant at the 95% threshold).

In Figure 3, we compare the evolution in time of the average cumulative infected patient-days under OPTI, COMBO, and OPTIDYN (see Figure A.7 in Appendix for percentiles). OPTI and COMBO tend to yield close results until approximately day 250. The performance of OPTIDYN, however, diverges from that of OPTI and COMBO as early as from day 100. Hence during the first 100 days, the information regarding the state of the epidemic is of little value – this period corresponds to the transient state departing from the initial treatment 0 steady state. However, after day 100, information has a significant value as it allows a substantial decrease in the number of infected individuals.

**Figure 3:**
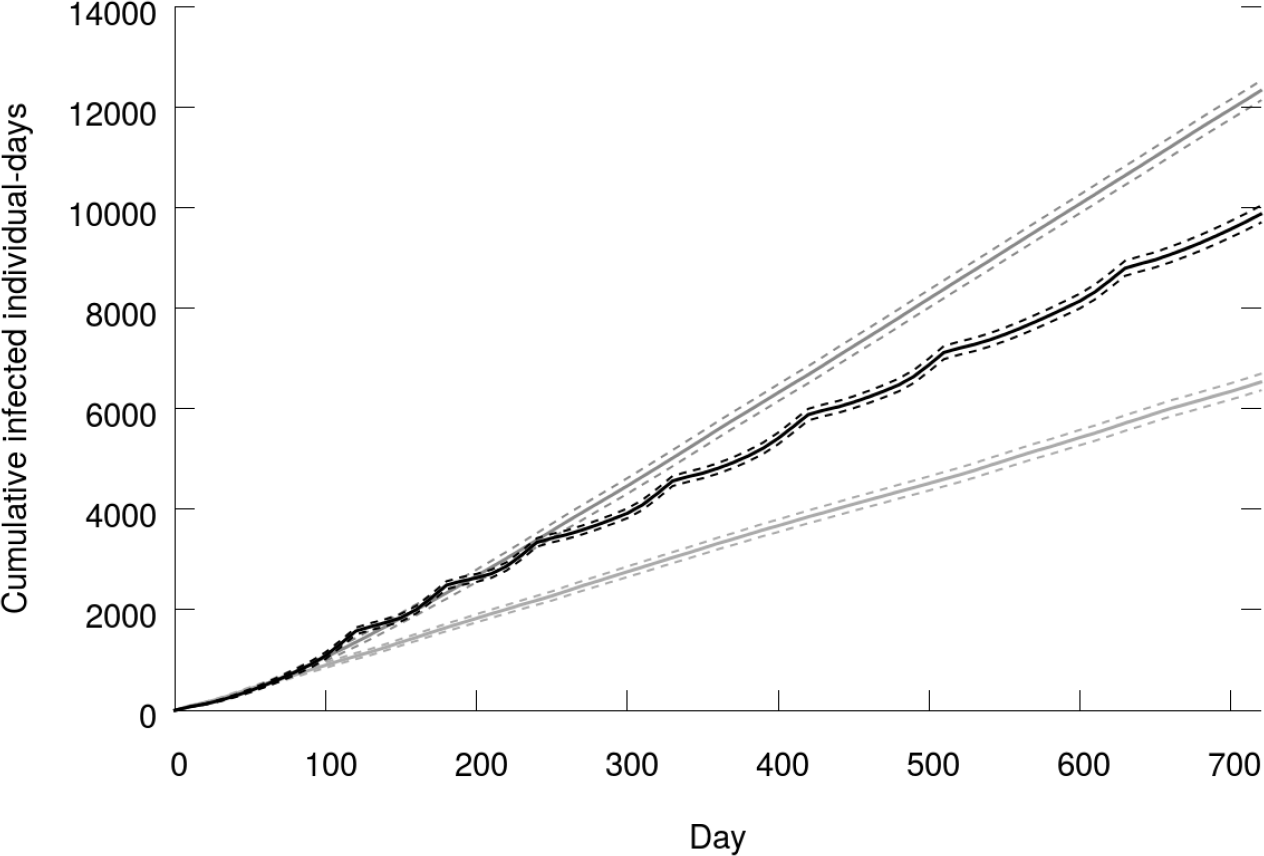
Average cumulative infected patient-days as a function of time (400 simulations). *Black:* OPTI. *Gray (above):* COMBO. *Gray (below):* OPTIDYN. *Dashed:* 95% CI.

We also ran simulations using our algorithm with Δ*h* = 30 days. We obtained an average of 7,378.65 (95% CI: 7,196.13 – 7,561.17) cumulative infected patient-days over two years. Thus, starting from optimization without surveillance (OPTI), implementing surveillance every 30 days and using this information to decide on empirical therapies allows a 23.4% decrease in average cumulative infected patient-days over two years. Doubling surveillance frequency to Δ*h* = 15 days yields a further 11.5% decrease. With Δ*h* = 5 days, we obtained 8,639.12 (95% CI: 8,420.17 – 8,858.06) cumulative infected patient-days over two years. In this case, the stochastic noise is too important to have better results than with Δ*h* = 15 days. Improving those results would require larger computational power.

In a health care facility, our optimization algorithm can be implemented *online* by running tests to get the number of patients with each health status every Δ*h* days, and then calling function STATETOTREATMENT to decide on an empirical treatment for the next Δ*h* days. The computation of an empirical treatment for the next Δ*h* days given the current state of the epidemic is tractable on a laptop computer. An alternative way to use the algorithm is to run simulations beforehand (just as we did to produce the results presented here), store the results, and use them to read directly the empirical treatment to be administered given the population with each health status, and the population evolution to be expected given that treatment. Doing so, it might prove necessary to resort to clustering techniques,^4^ which is left to later investigations.

## 4 Conclusion

We propose a solution to the problem of choosing an antimicrobial treatment to be administered empirically to the whole patient population in a health care facility. Our objective is to minimize the cumulative infected patient-days in the long run given that infecting organisms may evolve antimicrobial resistance. The empirical therapy is to be chosen periodically based on the count of patients with each health status at that moment, and on the population dynamics parameters of the health care facility and of the disease.

Previous studies have investigated empirical strategies based on screening test results. The great advantage of such strategies is that they make use of information available at no additional cost. However this approach typically only allows to compare a number of specific strategies. Also, these strategies are adaptive in that they specify an empirical treatment at each time without taking account of the subsequent spread of the disease.

To some extent, it could be said that practical considerations were the starting point of these previous studies. Here we looked at the problem the other way around by striving for near optimal empirical therapies first by using flexible methods that will allow to take account of any practical constraint in the future. To do this, we used a variant of the Monte-Carlo tree search algorithm, a method first devised in the field of artificial intelligence.

In our simulations, the method presented in this article allowed a 47.0% reduction in the average cumulative infected patient-days over two years compared to the combination therapy (a strategy widely used in the clinic). Compared to an optimization method that does not use periodic surveillance data, the average cumulative infected patient-days is reduced by 32.2%.

Finally, our algorithm can be used to produce data amenable to quantitative analysis. We believe that this calls for further investigation as it could allow to set up simple and practical decision rules to solve the complex problem of choosing empirical treatments in a health care facility.

## A Additional figures

**Figure A.1:**
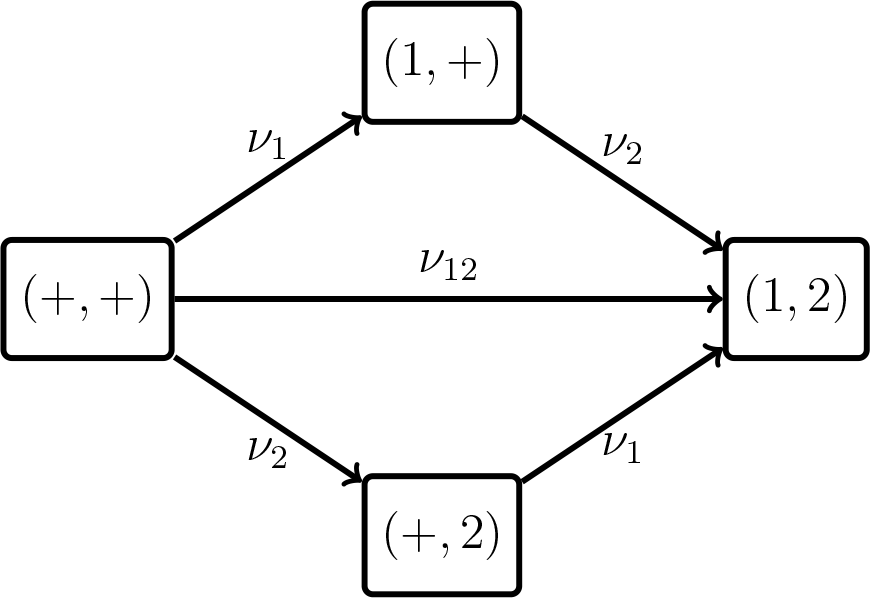
The four genotypes included in our model with mutation rates. The arrows represent transitions between genotypes. The two considered genes are separated by a comma. ‘+’ represents a wild-type allele. ‘1’ and ‘2’ represent alleles giving resistance to drugs 1 and 2 respectively. (+, +): wild-type susceptible. (1, +): 1-resistance only. (+, 2): 2-resistance only. (1, 2): resistance to both drugs.

**Figure A.2:**
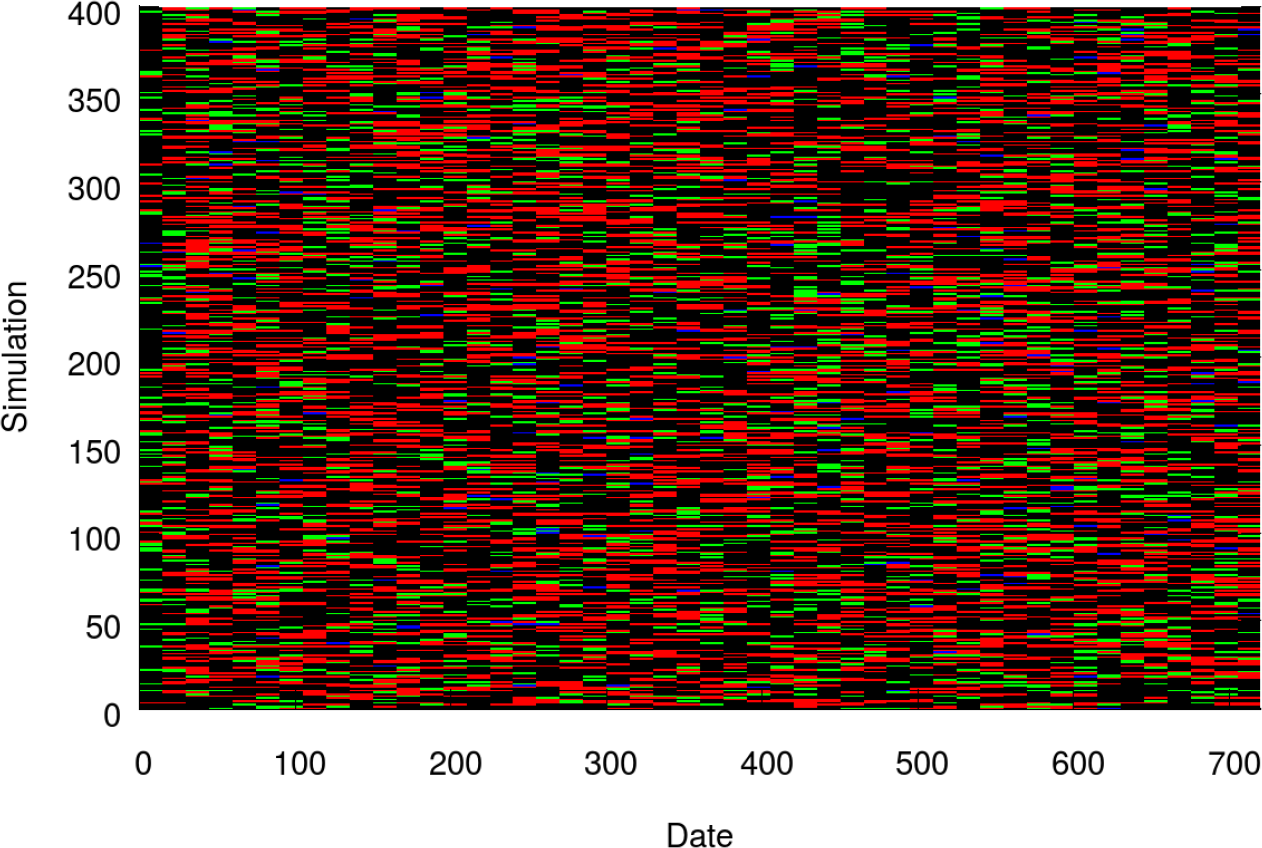
The 400 treatment regimens obtained with our optimization algorithm. *Red:* treatment 0. *Green:* treatment 1. *Blue:* treatment 2. *Black:* treatment 12.

**Figure A.3:**
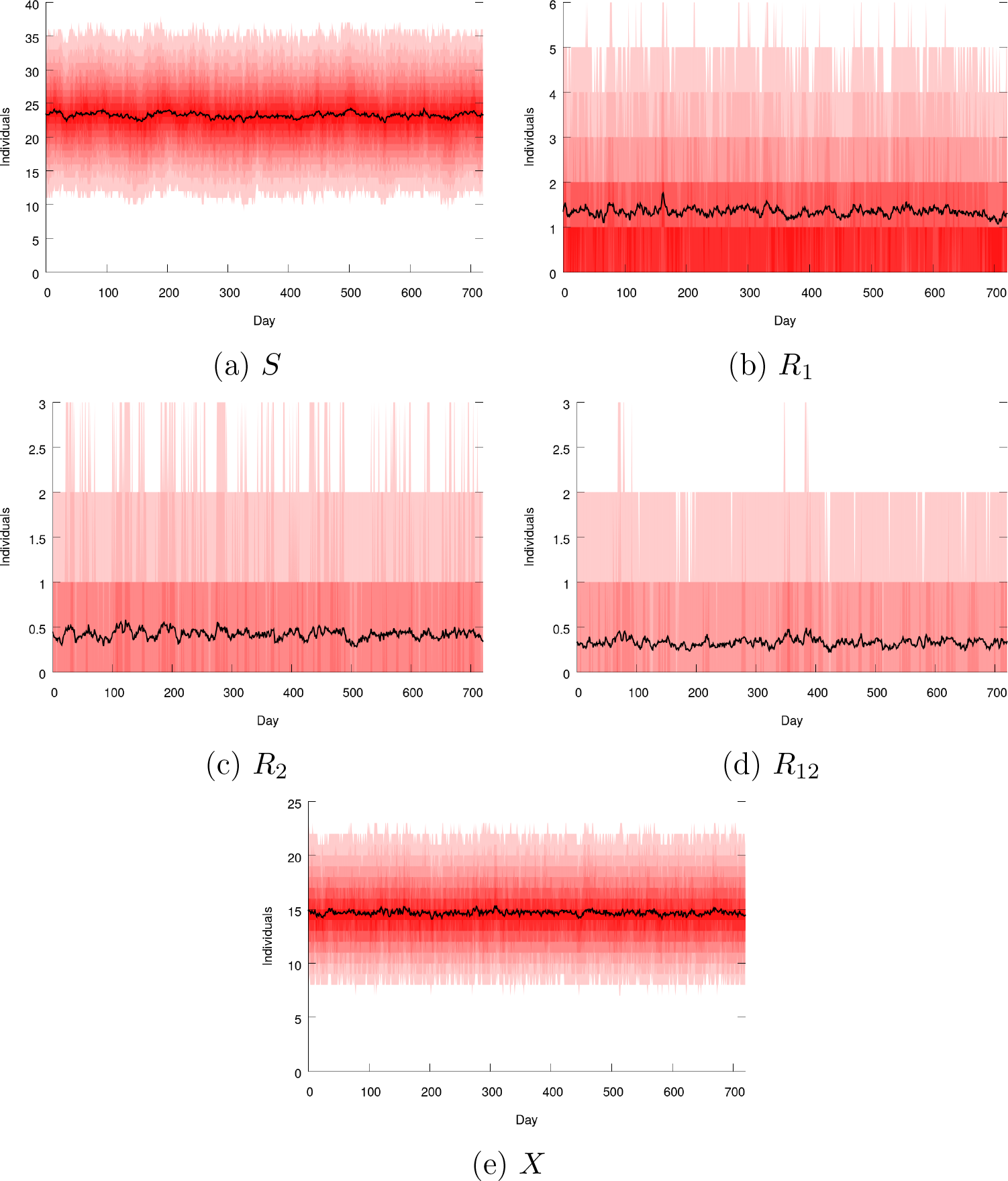
Number of patients in each compartment as a function of time when no treatment is used (NONE policy). *Black:* average over 400 simulations. *Shades of red: i*^th^ and (100 − *i*)^th^ percentiles for *i* from 5 to 50 by increments of 5.

**Figure A.4:**
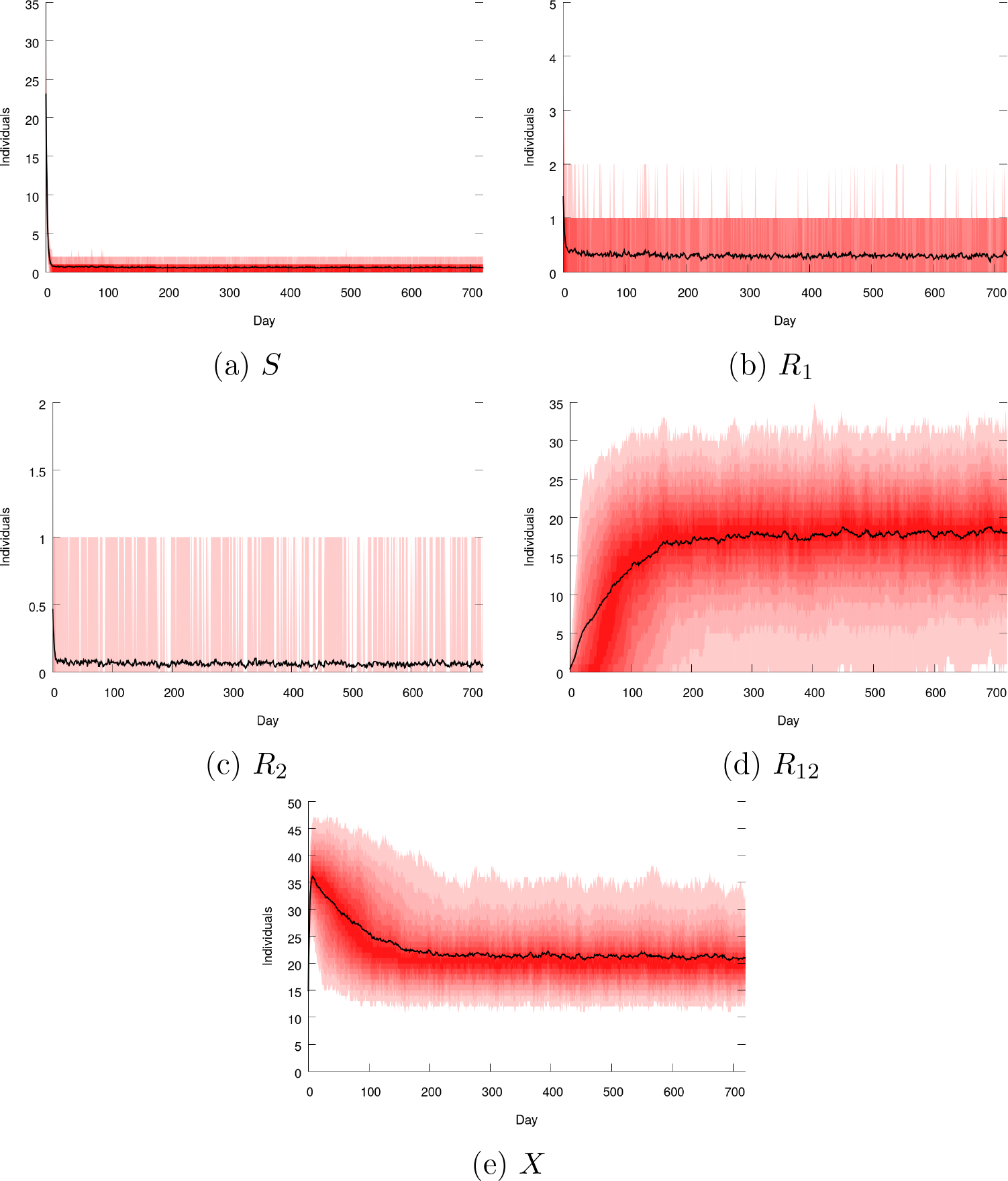
Number of patients in each compartment as a function of time when the COMBO policy is used. *Black:* average over 400 simulations. *Shades of red: i*^th^ and (100 − *i*)^th^ percentiles for *i* from 5 to 50 by increments of 5.

**Figure A.5:**
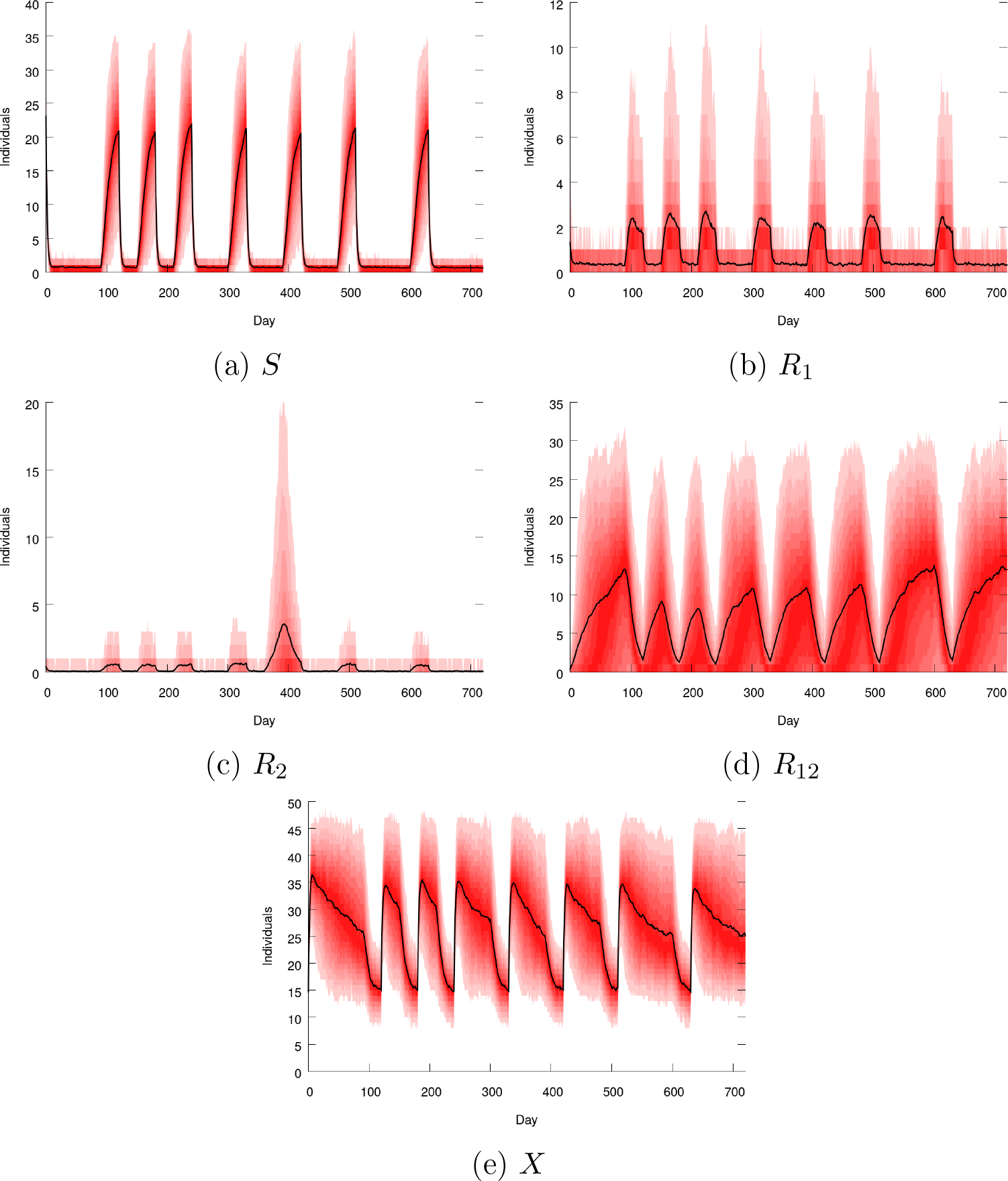
Number of patients in each compartment as a function of time when the OPTI policy is used. *Black:* average over 400 simulations. *Shades of red: i*^th^ and (100 − *i*)^th^ percentiles for *i* from 5 to 50 by increments of 5.

**Figure A.6:**
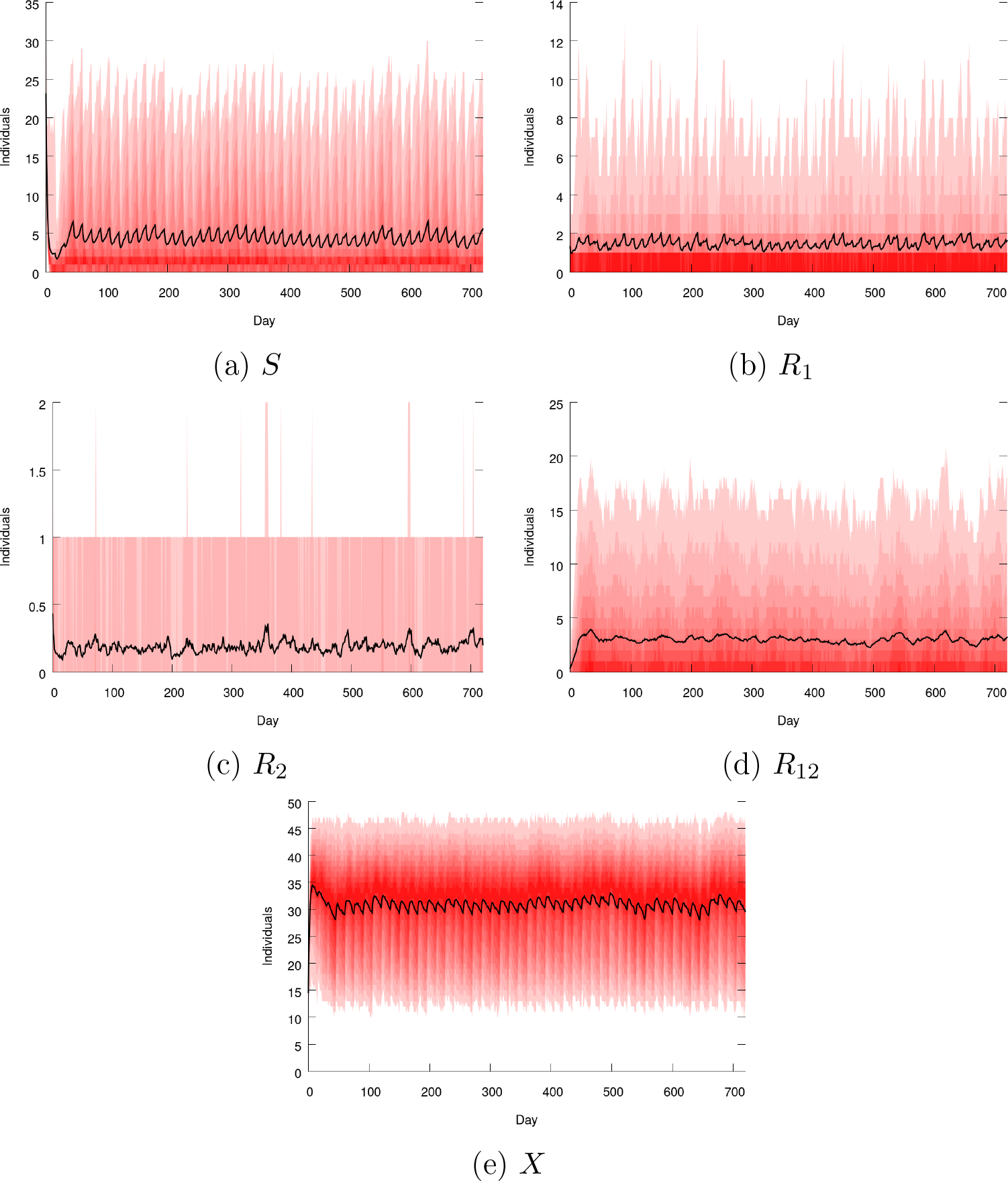
Number of patients in each compartment as a function of time when the OPTIDYN policy is used. *Black:* average over 400 simulations. *Shades of red: i*^th^ and (100 − *i*)^th^ percentiles for *i* from 5 to 50 by increments of 5.

**Figure A.7:**
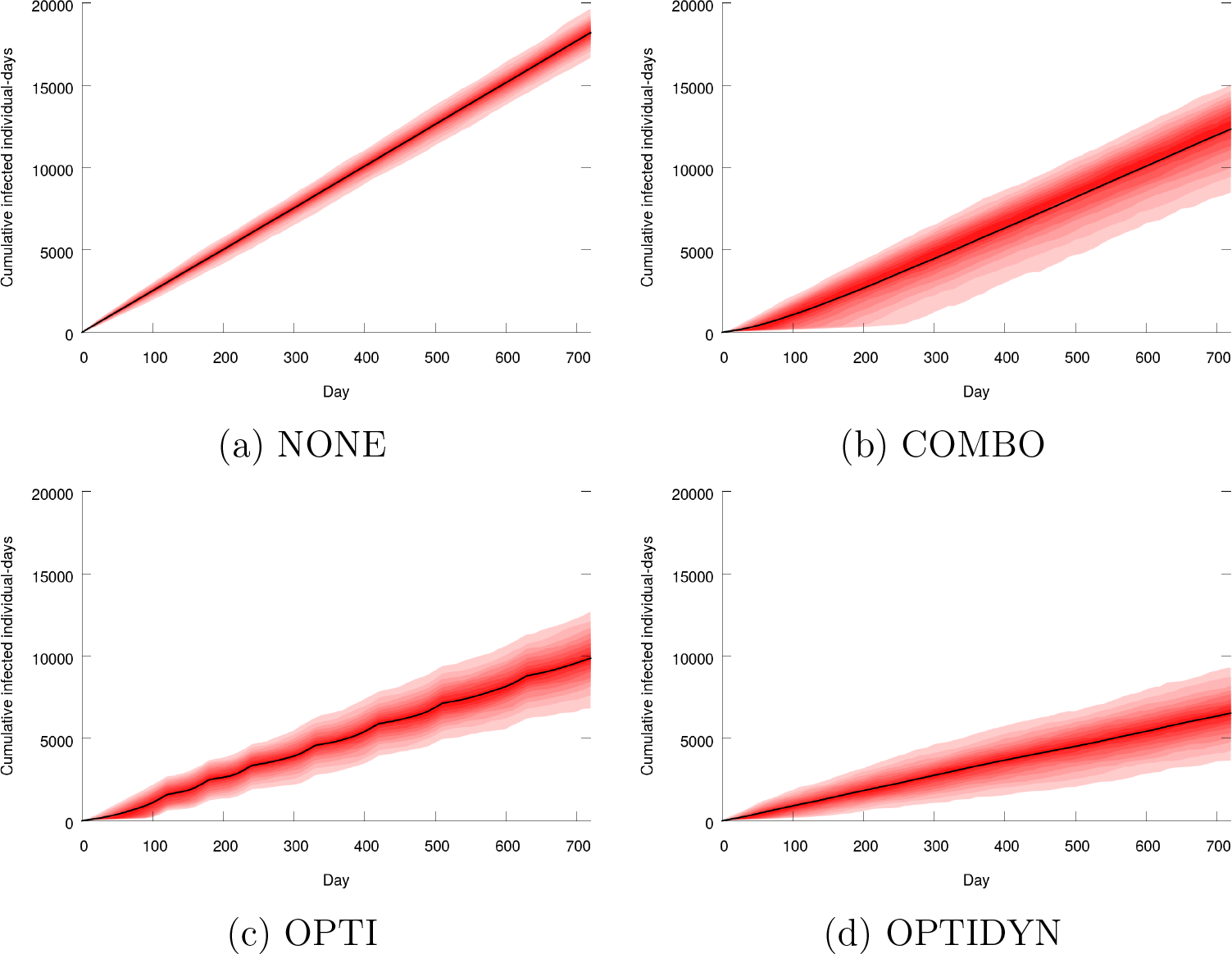
Cumulative infected patient-days as a function time when the NONE, COMBO, OPTI and OPTIDYN policies are used. *Black:* average over 400 simulations. *Shades of red: i*^th^ and (100 − *i*)^th^ percentiles for *i* from 5 to 50 by increments of 5.

**Figure A.8:**
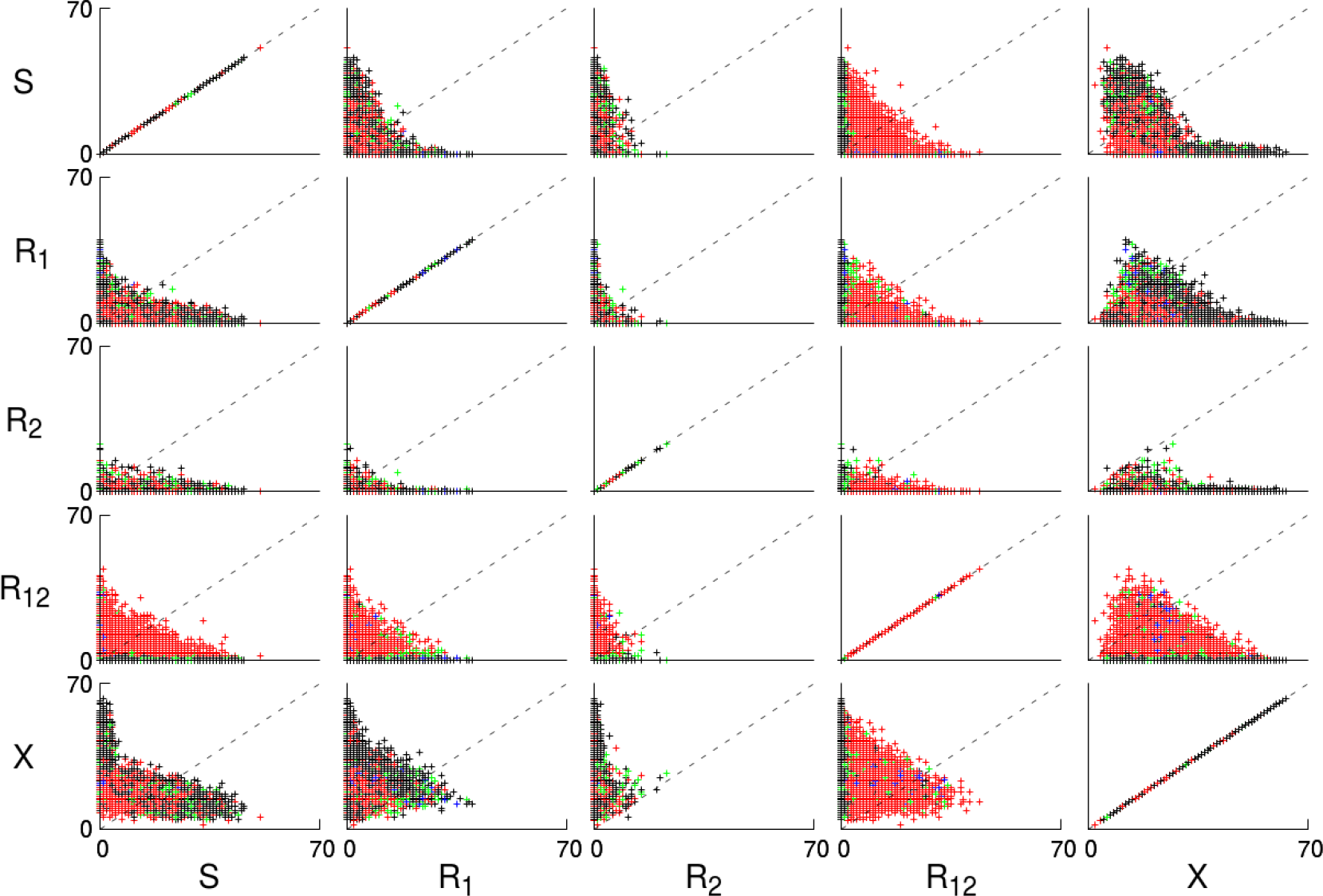
Chosen treatment as a function of the observed population in pairs of compartments. *Crosses:* function StateToTreatment calls. *Red:* treatment 0. *Green:* treatment 1. *Blue:* treatment 2. *Black:* treatment 12. *Dashed:* y=x lines.

## Footnotes

1 But not always, see [19].

2 Our definition of empirical therapy is therefore broader than the usual definition. We include prophy-lactic treatments in what we call empirical therapies for the sake of concision.

3 See Figures A.3–A.6 in Appendix for the evolution in time of the population in each compartment under different policies.

4 We illustrate the complexity of the output data of our model in Figure A.8 in Appendix.

## References

[1] Dan I. Andersson and Diarmaid Hughes. Antibiotic resistance and its cost: is it possible to reverse resistance? Nature Reviews Microbiology, 8(4):260, 2010.

[2] Robert Eric Beardmore, Rafael Peña-Miller, Fabio Gori, and Jonathan Iredell. Antibiotic cycling and antibiotic mixing: which one best mitigates antibiotic resistance? Molecular biology and evolution, 34(4):802–817, 2017.

[3] Carl T Bergstrom, Monique Lo, and Marc Lipsitch. Ecological theory suggests that antimicrobial cycling will not reduce antimicrobial resistance in hospitals. Proceedings of the National Academy of Sciences, 101(36):13285–13290, 2004.

[4] Sebastian Bonhoeffer, Pia Abel zur Wiesch, and Roger D Kouyos. Rotating antibiotics does not minimize selection for resistance. Math Biosci Eng, 7(4):919–22, 2010.

[5] Alessandro Cassini, Liselotte Diaz Högberg, Diamantis Plachouras, Annalisa Quattrocchi, Ana Hoxha, Gunnar Skov Simonsen, Mélanie Colomb-Cotinat, Mirjam E Kretzschmar, Brecht Devleesschauwer, Michele Cecchini, et al. Attributable deaths and disability-adjusted life-years caused by infections with antibiotic-resistant bacteria in the eu and the european economic area in 2015: a population-level modelling analysis. The Lancet Infectious Diseases, 19(1):56–66, 2019.

[6] Nicolas Houy and Julien Flaig. Optimal dynamic empirical therapy in a health care facility: an artificial intelligence approach. bioRxiv, 2019. Available at: https://doi.org/10.1101/603464.

[7] Nicolas Houy and François Le Grand. Optimizing immune cell therapies with artificial intelligence. Journal of Theoretical Biology, 461:34–40, jan 2019.

[8] Nicolas Houy and François Le Grand. Optimal dynamic regimens with artificial intelligence: The case of temozolomide. PLOS ONE, 13(6):1–15, 06 2018.

[9] Annette Jepson. Microbiology and infection control. In Carlos M H Gómez, editor, Clinical Intensive Care Medicine, chapter 10. Imperial College Press, 2014.

[10] Roger D. Kouyos, Pia Abel zur Wiesch, and Sebastian Bonhoeffer. Informed switching strongly decreases the prevalence of antibiotic resistance in hospital wards. PLoS computational biology, 7(3):e1001094, 2011.

[11] Bruce R Levin and Marc JM Bonten. Cycling antibiotics may not be good for your health. Proceedings of the National Academy of Sciences, 101(36):13101–13102, 2004.

[12] Marc Lipsitch and Matthew H. Samore. Antimicrobial use and antimicrobial resistance: a population perspective. Emerging infectious diseases, 8(4):347–354, 2002.

[13] Anita H. Melnyk, Alex Wong, and Rees Kassen. The fitness costs of antibiotic resistance mutations. Evolutionary applications, 8(3):273–283, 2015.

[14] Michael S. Niederman. Is “crop rotation” of antibiotics the solution to a “resistant” problem in the icu? American Journal of Respiratory and Critical Care Medicine, 156(4):1029–1031, 1997.

[15] R Peña-Miller and Robert Beardmore. Rotating antibiotics selects optimally against antibiotic resistance, in theory. Mathematical Biosciences & Engineering, 7(3):pp. 527–552, 2010.

[16] Rafael Peña-Miller and Robert Beardmore. Antibiotic cycling versus mixing: the difficulty of using mathematical models to definitively quantify their relative merits. Mathematical Biosciences & Engineering, 7(4):923–933, 2010.

[17] D. E. Ramsay, J. Invik, S. L. Checkley, S. P. Gow, N. D. Osgood, and C. L. Waldner. Application of dynamic modelling techniques to the problem of antibacterial use and resistance: a scoping review. Epidemiology & Infection, 146(16):2014–2027, 2018.

[18] Timothy C. Reluga. Simple models of antibiotic cycling. Mathematical medicine and biology: a journal of the IMA, 22(2):187–208, 2005.

[19] Damien Roux, Olga Danilchanka, Thomas Guillard, Vincent Cattoir, Hugues Aschard, Yang Fu, Francois Angoulvant, Jonathan Messika, Jean-Damien Ricard, John J Mekalanos, et al. Fitness cost of antibiotic susceptibility during bacterial infection. Science translational medicine, 7(297):297ra114–297ra114, 2015.

